# Beyond where: When and how brain stimulation drives state transitions

**DOI:** 10.64898/2026.03.17.712405

**Authors:** Irene Acero-Pousa, Leonardo Bonetti, Mattia Rosso, Yonatan Sanz-Perl, Peter Vuust, Morten L. Kringelbach, Gustavo Deco

## Abstract

Brain stimulation is increasingly used to treat neurological and psychiatric conditions, but stimulation sites and timing are typically chosen based on a priori assumptions from prior studies, not individualised mechanisms. Here, we investigate why some brain regions respond more to stimulation, when they respond best, and whether optimal targets for brain-state transitions are defined by functional anatomy (e.g. an auditory region in a resting-to-listening task transition), by system dynamics, or a combination of both. To do so, we build subject-specific whole-brain Hopf models fitted to MEG data, separating signals into five canonical frequency bands. We find that regions with a smaller oscillation radius and greater temporal variability respond more strongly to stimulation. Moreover, responsiveness was highly dependent on state and frequency band, with slower frequencies exhibiting phase-driven responsiveness, while faster ones depending more on network synchrony. Importantly, optimal nodes for transitioning between brain states are not defined by functional labels but rather by their dynamical regime. These results emphasise the value of mechanistic, personalised models for identifying when and where to stimulate the brain, and may help inform more precise brain-stimulation strategies.

## 1 Introduction

Brain stimulation is increasingly used as a therapeutic intervention across neurological and psychiatric disorders, either as a primary treatment or in combination with pharmacological approaches to enhance efficacy. Clinical benefits have been demonstrated in Parkinson’s disease [1], obsessive–compulsive disorder and major depression, among others, for which several stimulation-based interventions are now in clinical use [2]. However, current stimulation protocols remain largely empirical, with parameters that are typically chosen based on what works best on average at the group level, and with limited consideration of between-subject variability [3, 4, 5], even though brain activity can differ substantially between patients with the same diagnosis [6]. Variability also arises within individuals: not all stimulation sites or time points elicit the same response [3, 7, 8], and the effect of a given protocol depends on the brain state (e.g. awake vs. anaesthetised) at the moment of stimulation [9, 10, 11].

This variability calls for mechanistic frameworks that can relate individual brain activity to its response to perturbation. Whole-brain computational models provide such a framework [12, 13, 14]. By combining empirically derived structural and functional connectivity with biophysically inspired local dynamics, these models capture selected spatiotemporal features at the level of each individual. Once tuned to a person’s empirical data, the model can be perturbed *in silico* to probe the causal impact of stimulating different regions, time points or intensities, and to quantify how stimulation reshapes whole-brain dynamics [15, 16, 17]. This allows systematic exploration of the “where, when and how” of stimulation in a way that would be infeasible or unethical to test exhaustively *in vivo* [18].

Previous work using whole-brain modeling has focused on the spatial dimension of this problem. Perturbation studies have shown that, in these models, the impact of local stimulation depends on a region’s embedding in the network and its position along cortical hierarchies: structurally central regions often tend to have slower, more stable dynamics, whereas peripheral regions show faster, higher-variance activity and larger changes in functional connectivity when perturbed [19, 20, 21]. In line with this view, our recent Inception work on disorders of consciousness found that regions with higher effective-connectivity degree were less effective at driving the desired transitions between brain states, suggesting that excessive integration can make nodes harder to steer [22]. Complementary work combining intracranial stimulation with whole-brain modeling has revealed gradients of excitability and recurrence across cortex, with higher-order association areas exhibiting stronger and more prolonged network responses due to recurrent feedback from the rest of the brain [16]. Yet it remains unclear whether the most effective stimulation targets for a desired state transition are primarily defined by their anatomical role in the target network, or instead by more generic properties that make certain nodes intrinsically easier to steer. Moreover, a node’s responsiveness to stimulation (how much it responds when perturbed) need not coincide with its effectiveness at driving a specific transition in state space.

By contrast, the temporal dimension of when to stimulate (i.e., how the effect of a perturbation depends on its precise timing relative to ongoing brain activity) has been explored extensively *in vivo* and in circuit-level models, where phase- or state-dependent stimulation can produce different outcomes [23, 24, 25]. At the whole-brain scale, however, what is still missing is a mechanistic, personalised map to understand which features of the ongoing dynamics predict when a given node is maximally responsive, and how these rules generalise across regions and frequency bands.

In this study, we pursue two complementary goals. First, we extend previous work by asking not only where but also when to perturb, and we use machine learning to predict, for each region, the stimulation time points that are expected to be maximally responsive. Second, we ask whether, for a given brain-state transition, the optimal stimulation sites are primarily determined by target-relevant systems (e.g. auditory cortex for a rest-to-listen transition) or by more general spatial and dynamical response properties of the network. To address these questions, we analyse magnetoencephalography (MEG) data from 28 participants recorded during rest and during a passive listening task. For each participant, state (i.e., resting vs. listening) and canonical frequency band, we fit a personalised whole-brain model. We then systematically perturb the model at each brain region at different time points to characterise where and when stimulation elicits the largest responses (i.e., the connectivity after the stimulation is most different from before) . Finally, we identify the regions whose stimulation in the resting-state model is most effective at driving the dynamics toward the listening state (i.e., the connectivity after the stimulation is most similar to the listening connectivity), and test whether these optimal targets coincide with auditory regions or instead reflect generic spatial and temporal response mechanisms.

In summary, we show that stimulation effects in personalised whole-brain models are strongly band- and state-dependent, and that a node’s responsiveness to stimulation can be explained by its dynamical regime (amplitude of the oscillation). In terms of timing, we demonstrate that the instantaneous local phase and network synchrony predict how strongly a node responds at a given time, and can be used with machine learning to select stimulation time points that consistently outperform random timing. For the rest-to-listening transition, the nodes that best drive the state change are not confined to auditory cortex, but instead share the same dynamical fingerprints identified in the responsiveness analyses. Together, these results point to a dynamical, rather than purely anatomical, principle for where and when to perturb the brain to induce desired state transitions.

## 2 Results

In this paper, our aim was twofold. First, we asked which static and time-resolved mechanisms determine how strongly a node responds to an external stimulation. Second, we asked whether the nodes that are most effective for driving a transition toward a different brain state are mainly target-dependent or instead determined by more general mechanisms.

To address these questions, we followed the steps summarised in **Figure 1**. For each state (resting and listening), we first divided the MEG time series of each participant into the five canonical frequency bands (**Figure 1a, Figure 1b**). For each participant and frequency band, we built personalised whole-brain models and estimated the corresponding Generative Effective Connectivity (GEC; see **Methods**; **Figure 1c**). The fitting accuracy of these models is described in **Table 1**. We then stimulated each brain region independently at different time points (evenly spaced intervals). For each stimulation, we computed the functional connectivity (FC) of the stimulated data and quantified the response as the L2 distance between this FC and the FC obtained before any stimulation, restricting the distance to the row of connections of the stimulated node. This yielded, for every node and frequency band, a distribution of responses across stimulated time points (**Figure 1d**). We then examined how these responses relate to static (time-aggregated summaries that are constant across stimulation onsets, e.g., structural connectivity strength) and time-resolved (features computed at the exact onset, e.g., phase) mechanisms, and finally tested whether the optimal nodes for the transition between the resting and listening states were auditory (target-state driven) or instead explained by the same static mechanisms.

**Table 1:** Model fitting accuracy by frequency band and condition. Values show mean *±* standard deviation across participants of the Pearson correlation between empirical and model-predicted connectivity matrices. *FC* denotes the zero-lag functional connectivity (correlation) matrix. *FS*(*τ*) denotes the normalized time-shifted covariance matrix used for optimization, computed at *τ* = 3 samples for delta–alpha and *τ* = 2 samples for beta–gamma.

| **Band** | **REST $FC$** | **REST $FS(\tau)$** | **LISTEN $FC$** | **LISTEN $FS(\tau)$** |
| --- | --- | --- | --- | --- |
| Delta | $0.615 \pm 0.020$ | $0.669 \pm 0.020$ | $0.605 \pm 0.018$ | $0.661 \pm 0.024$ |
| Theta | $0.599 \pm 0.023$ | $0.640 \pm 0.032$ | $0.602 \pm 0.028$ | $0.647 \pm 0.035$ |
| Alpha | $0.595 \pm 0.014$ | $0.607 \pm 0.037$ | $0.593 \pm 0.009$ | $0.563 \pm 0.108$ |
| Beta | $0.638 \pm 0.038$ | $0.631 \pm 0.042$ | $0.609 \pm 0.022$ | $0.606 \pm 0.025$ |
| Gamma | $0.624 \pm 0.022$ | $0.662 \pm 0.029$ | $0.609 \pm 0.016$ | $0.649 \pm 0.021$ |

**Figure 1:**
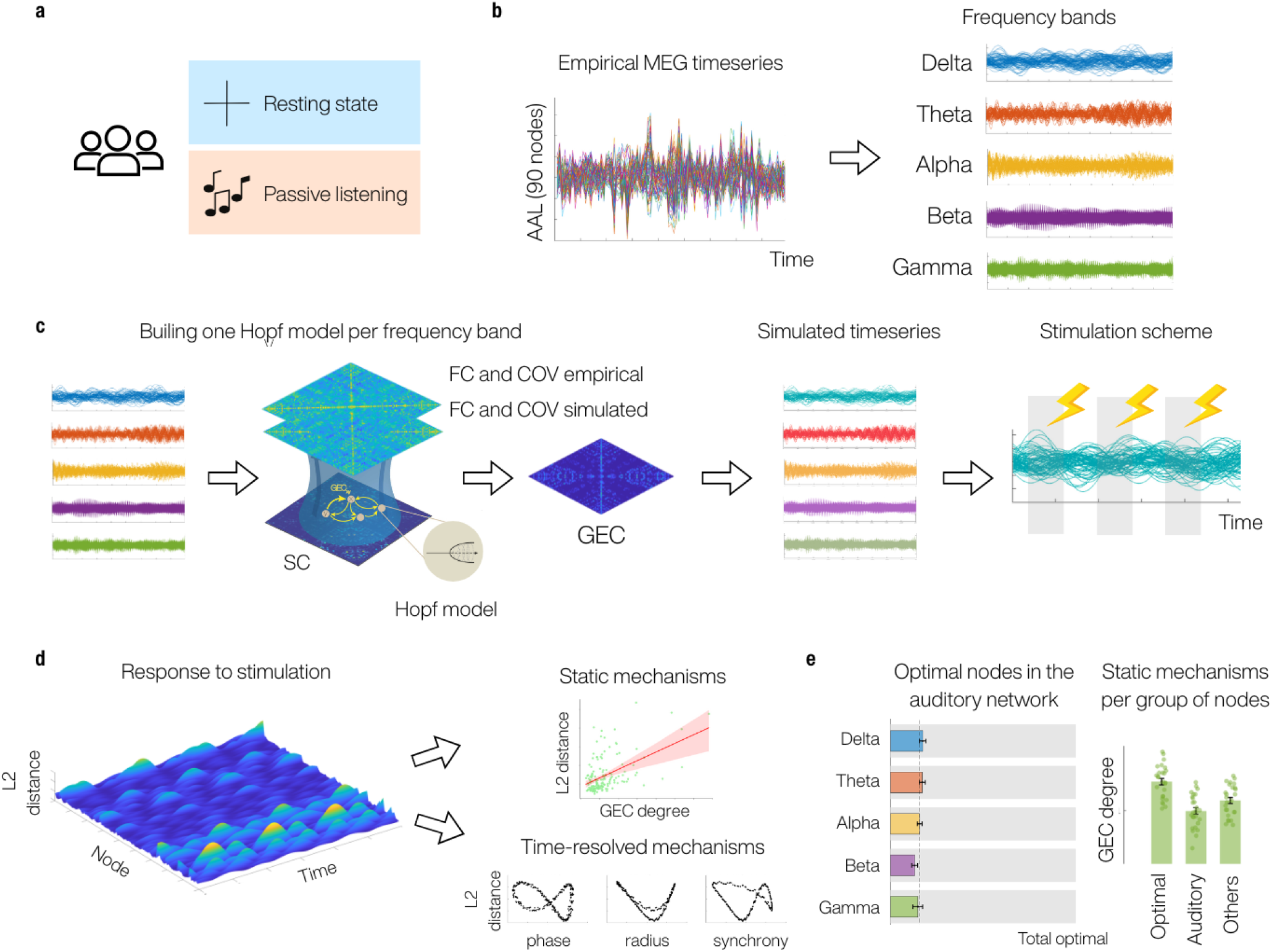
Modelling pipeline to characterise static and dynamic mechanisms of node responses and optimal stimulation targets. **(a)** Dataset of 28 participants scanned during resting state and a passive listening task. **(b)** Source-reconstructed MEG time series were parcellated (AAL90) and split into five canonical frequency bands (delta–gamma). **(c)** For each participant and band, we built a Hopf whole-brain model whose parameters were optimised to match empirical FC and covariance, yielding a subject- and band-specific generative effective connectivity (GEC). Using this GEC, we simulated band-limited time series and applied a sinusoidal drive to individual nodes at multiple stimulation time points. **(d)** For each node and time point, the response to stimulation was quantified as the L2 distance between the pre-stimulus FC and the post-stimulus FC row of the stimulated node. We then related response magnitude and variability to static node properties (in-degree, out-degree, eigenvector centrality, mean radius and radius standard deviation) and to a set of 12 time-resolved features capturing local phase, radius, input and network synchrony. **(e)** Finally, we used the same framework to identify, for each band, the nodes that best drive the transition from rest to listening and tested whether these optimal targets are primarily determined by belonging to the auditory network or by their static dynamical regime.

### 2.1 Response magnitude and variability over time is band-specific and state-specific

We first explored whether response magnitude differs across frequency bands, and whether this pattern is consistent between the resting state and the listening task. For each band, response magnitude was computed as the average over all stimulated time points for each brain region, and then averaged across regions, obtaining one value per participant. In the resting state, we observed a significant decrease in response magnitude from slow to fast bands, with delta showing the largest responses (**Figure 2b**). However, this pattern did not hold during the listening task, where the decrease was preserved for all bands except alpha, which showed the highest responses (**Figure 2c**). This pattern is summarised in **Figure 2d**, which directly compares both states. Magnitude was significantly higher during rest than listening for delta (p *<* 0.001), whereas alpha showed significantly higher magnitude during listening than rest (p *<* 0.001), with no clear state differences in theta, beta or gamma (p *>* 0.05).

**Figure 2:**
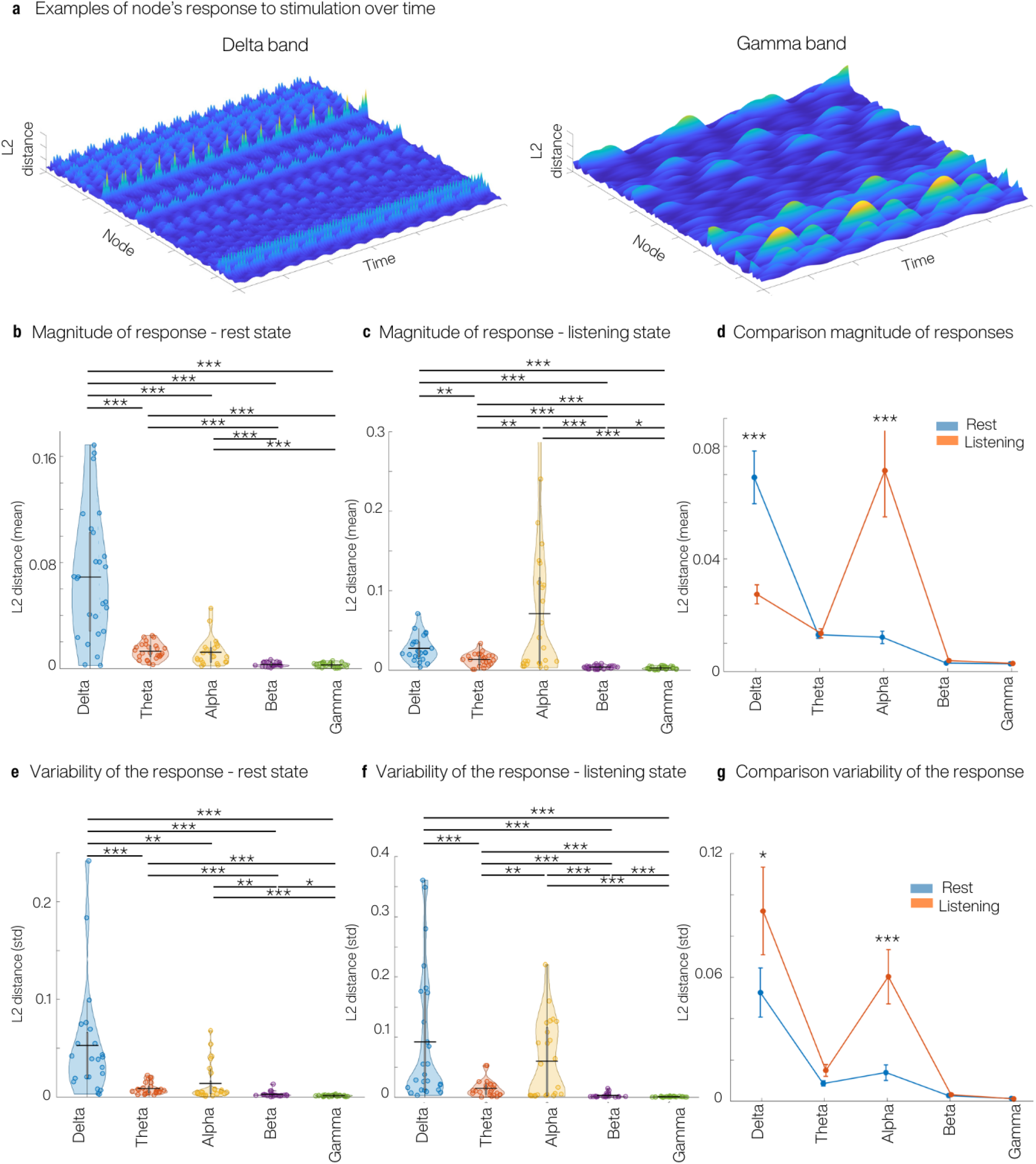
Response magnitude and variability to external stimulation are band- and state-dependent. **(a)** Example node-wise and time-resolved responses in delta and gamma bands for one participant. **(b)** Magnitude of the response (mean L2 distance) across frequency bands in the resting state. **(c)** Magnitude of the response across bands in the listening state. **(d)** Comparison of response magnitude between states: rest shows a clear maximum in delta, whereas listening shows a selective increase in alpha. The rest of the bands show similar magnitude of responses. **(e)** Variability of the response over time (standard deviation of L2 distance) across bands in the resting state. **(f)** Variability of the response over time across bands in the listening state. **(g)** Comparison of response variability between states: variability is highest in delta at rest and increases selectively in alpha during listening, while the remaining bands show smaller differences. All p-values are FDR corrected, * indicates p *<* 0.05, ** indicates p *<* 0.01 and *** indicates p *<* 0.001

We then repeated the analysis using response variability instead of magnitude. Variability was quantified as the standard deviation across stimulated time points for each brain region, and then averaged across regions. In the resting state, we again observed a strong band-dependence, with delta showing the largest variability, which progressively decreased toward faster bands, with alpha remaining slightly higher than theta and beta (**Figure 2e**). In the listening state, the profile changed. Delta remained highly variable, but alpha also showed a marked increase in variability and clearly stood out from the intermediate bands (**Figure 2f**). When directly comparing states, variability was significantly higher during listening than rest for delta (p *<* 0.05) and especially for alpha (p *<* 0.001), whereas theta, beta and gamma showed no clear state differences (p *>* 0.05, **Figure 2g**).

### 2.2 Static mechanisms of response magnitude and variability over time

Next, we asked which static mechanisms explain why some nodes respond more than others, and why some have a more variable response over time. For each band, state and participant, we correlated node-wise response magnitude/variability with five static properties: in-degree, out-degree and eigenvector centrality from the GEC, and the mean and standard deviation (std) of the baseline radius time series. In-degree and out-degree quantify how many directed connections a node receives and sends, and eigenvector centrality quantifies its embedding in the network. Mean radius and std radius quantify the node’s oscillation amplitude and the temporal variability of this amplitude, respectively (see **Methods**). This gave one Spearman correlation value per subject, metric and band (**Figure 3, Figure S1** and **Tables 2**, **3**).

**Table 2:** Static mechanisms of stimulation effect by frequency band and state. Values show mean *±* standard deviation across participants (*n* = 28) of the Spearman correlation (*ρ*) between node-wise static metrics (in-degree, out-degree, eigencentrality, mean radius, and radius standard deviation) and the node-wise stimulation effect. For each participant and band, the stimulation effect per node is defined as the temporal mean of the effect time course, *ρ* is computed across nodes and then summarised across participants.

| Band | REST |  |  |  |  | LISTEN |  |  |  |  |
| --- | --- | --- | --- | --- | --- | --- | --- | --- | --- | --- |
|  | In-deg | Out-deg | Eigen | MeanR | StdR | In-deg | Out-deg | Eigen | MeanR | StdR |
| Delta | -0.230 ± 0.170 | -0.230 ± 0.171 | -0.133 ± 0.143 | -0.746 ± 0.109 | 0.778 ± 0.122 | -0.080 ± 0.199 | -0.081 ± 0.199 | -0.024 ± 0.182 | -0.346 ± 0.188 | 0.330 ± 0.207 |
| Theta | 0.082 ± 0.168 | 0.075 ± 0.168 | 0.126 ± 0.156 | -0.421 ± 0.159 | 0.394 ± 0.170 | 0.040 ± 0.188 | 0.037 ± 0.189 | 0.069 ± 0.184 | -0.165 ± 0.152 | 0.143 ± 0.151 |
| Alpha | -0.023 ± 0.145 | -0.021 ± 0.147 | 0.015 ± 0.141 | -0.305 ± 0.128 | 0.281 ± 0.186 | 0.010 ± 0.111 | 0.009 ± 0.112 | 0.019 ± 0.129 | -0.073 ± 0.146 | 0.069 ± 0.128 |
| Beta | -0.209 ± 0.212 | -0.215 ± 0.221 | -0.038 ± 0.182 | -0.675 ± 0.316 | 0.702 ± 0.283 | -0.045 ± 0.155 | -0.056 ± 0.153 | 0.003 ± 0.136 | -0.247 ± 0.203 | 0.211 ± 0.207 |
| Gamma | -0.071 ± 0.226 | -0.074 ± 0.230 | 0.048 ± 0.195 | -0.747 ± 0.225 | 0.741 ± 0.229 | -0.028 ± 0.176 | -0.028 ± 0.179 | 0.036 ± 0.167 | -0.283 ± 0.219 | 0.243 ± 0.226 |

**Table 3:** Static mechanisms of the *temporal variability* of stimulation effect by frequency band and condition. Values show mean *±* standard deviation across participants (*n* = 28) of the Spearman correlation (*ρ*) between node-wise static metrics and the node-wise temporal standard deviation of the stimulation effect time course. For each participant and band, *ρ* is computed across nodes and then summarized across participants.

| Band | REST |  |  |  |  | LISTEN |  |  |  |  |
| --- | --- | --- | --- | --- | --- | --- | --- | --- | --- | --- |
|  | In-deg | Out-deg | Eigen | MeanR | StdR | In-deg | Out-deg | Eigen | MeanR | StdR |
| Delta | -0.182 ± 0.141 | -0.182 ± 0.141 | -0.123 ± 0.135 | -0.535 ± 0.134 | 0.581 ± 0.128 | -0.080 ± 0.188 | -0.081 ± 0.188 | -0.034 ± 0.171 | -0.308 ± 0.184 | 0.295 ± 0.200 |
| Theta | 0.075 ± 0.169 | 0.068 ± 0.169 | 0.115 ± 0.157 | -0.379 ± 0.165 | 0.355 ± 0.173 | 0.040 ± 0.179 | 0.039 ± 0.181 | 0.063 ± 0.182 | -0.143 ± 0.155 | 0.122 ± 0.154 |
| Alpha | -0.052 ± 0.146 | -0.051 ± 0.148 | -0.018 ± 0.149 | -0.237 ± 0.127 | 0.224 ± 0.170 | 0.006 ± 0.118 | 0.003 ± 0.120 | 0.017 ± 0.139 | -0.057 ± 0.130 | 0.066 ± 0.122 |
| Beta | -0.156 ± 0.264 | -0.166 ± 0.276 | -0.007 ± 0.202 | -0.621 ± 0.408 | 0.603 ± 0.450 | -0.054 ± 0.140 | -0.066 ± 0.137 | -0.006 ± 0.123 | -0.231 ± 0.205 | 0.207 ± 0.198 |
| Gamma | -0.099 ± 0.212 | -0.100 ± 0.216 | 0.025 ± 0.199 | -0.696 ± 0.324 | 0.695 ± 0.299 | -0.049 ± 0.177 | -0.054 ± 0.175 | 0.002 ± 0.165 | -0.239 ± 0.237 | 0.205 ± 0.241 |

**Figure 3:**
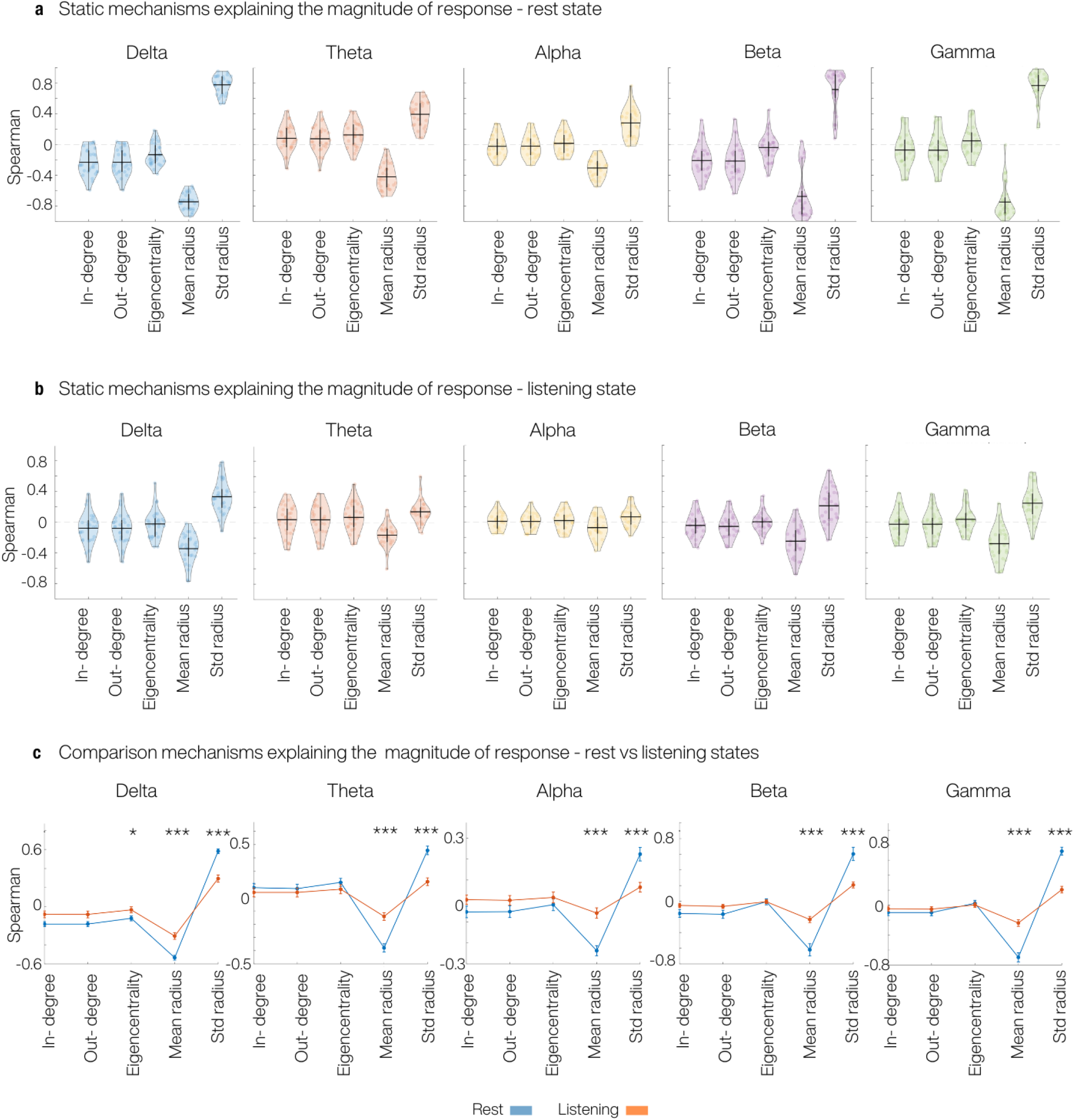
Node dynamics explains response magnitude after stimulation. **(a)** Distribution across participants of Spearman correlations between node-wise response magnitude and five static metrics (in-degree, out-degree, eigenvector centrality, mean baseline radius and baseline radius variability) for each frequency band in the resting state. **(b)** Same as in (a), but for listening state instead of the resting state. **(c)** Direct comparison of mechanisms between rest and listening for the response magnitude. Lines show the mean *±* SEM of the Spearman correlations across participants, separately for each band and metric, asterisks indicate significant rest–listening differences (FDR-corrected, * indicates p *<* 0.05 and *** indicates p *<* 0.001). Across bands, mean radius correlates negatively and std radius positively with the magnitude of the response, with systematically stronger effects at rest than during listening.

In the resting state, both response magnitude and variability were mainly explained by the dynamical properties of the nodes (**Figure 3a**, **Figure S1a**). Across all bands, mean radius tended to correlate negatively with response magnitude and variability, whereas std radius showed the strongest and most consistent positive correlations, particularly in the slower bands. In contrast, the structural metrics (in-degree, out-degree and eigenvector centrality) showed only modest and band-dependent correlations, often close to zero and sometimes changing sign across bands.

During the listening task, the qualitative pattern was similar but markedly weaker (**Figure 3b**, **Figure S1b**). The dynamical metrics remained the most informative: mean radius tended to correlate negatively and std radius positively with both response magnitude and variability, but the corresponding correlation coefficients were smaller than at rest. Structural metrics again contributed little, with narrow distributions centred near zero in most bands.

The direct comparison between states confirmed this difference (**Figure 3c**, **Figure S1c**). Overall, the same static mechanisms are relevant in both conditions, but they are significantly stronger at rest than during listening (p *<* 0.001 for mean and std radius across bands).

### 2.3 Time-specific mechanisms shaping a node’s response

We followed our analysis by asking whether the instantaneous dynamical state of the system constrains how strongly a node reacts to stimulation at a given time point. For each band, node and participant, we extracted a set of time-specific features at every stimulation time point: local phase (cosine and sine), local radius, magnitude and radial component of the input, global and local synchrony, global radius, and short-term histories of the global and local radius (**Figure 4a right**, see **Methods** for specific definitions). We then trained a random forest regression model that, based on all these features, predicted response magnitude at each time point. Model performance was quantified with out-of-bag *R*^2^ for each node. The average *R*^2^ was clearly above zero in all frequency bands, showing that the momentary dynamical state contains substantial information about how much a node will respond to stimulation (**Figure 4a left**). Importantly, these models also provided, for each brain region, a feature-importance profile indicating which time-specific factors were most informative for each of them.

**Figure 4:**
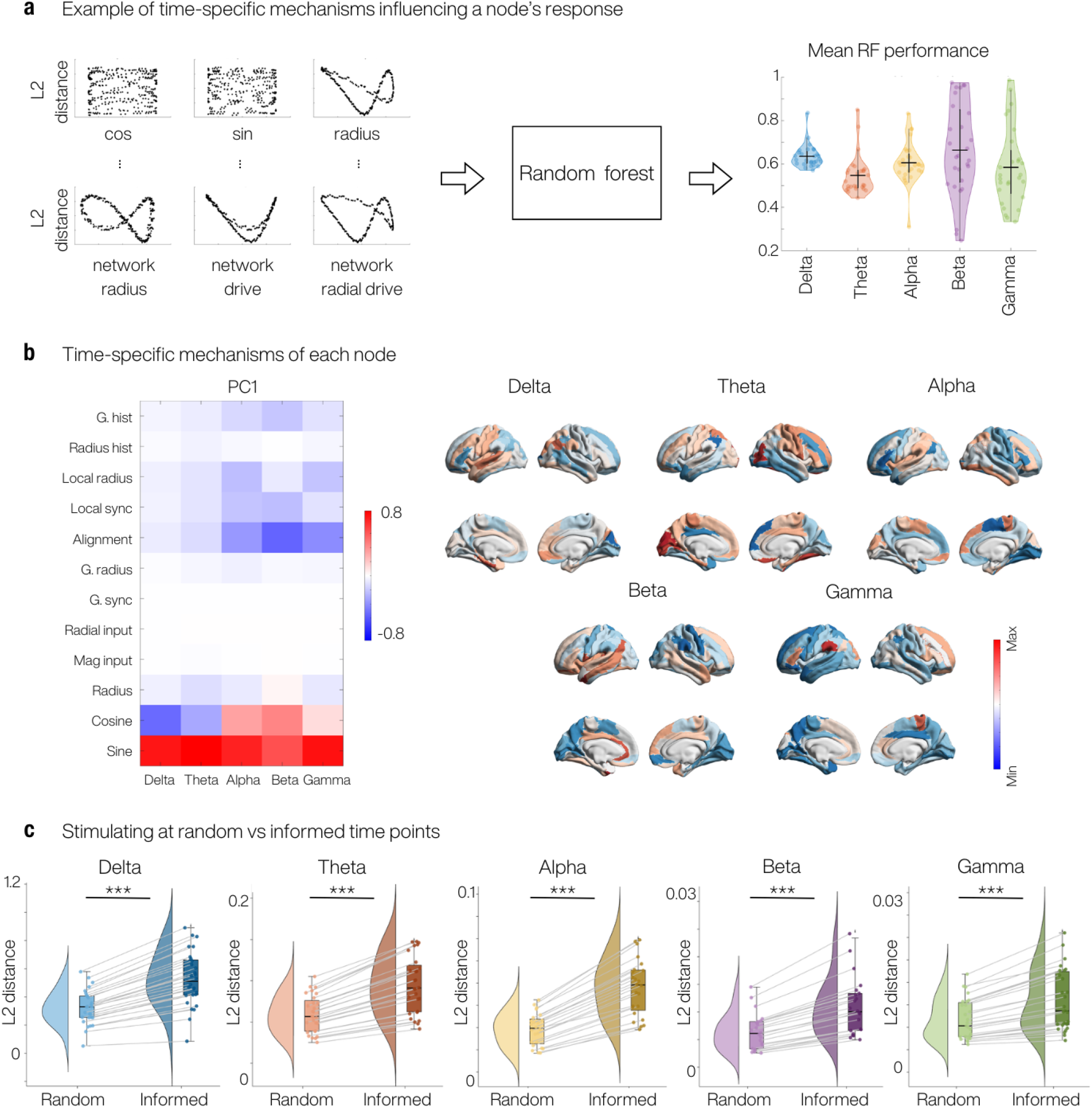
Time-specific dynamical states explain node responses and can be used to optimise stimulation timing. **(a)** Schematic of the random forest regression approach. For each stimulation time point we extract a set of time-specific features (local phase, local radius, input, global and local synchrony and their histories) and train a model to predict the response magnitude. Violin plots show the distribution across participants of out-of-bag *R*^2^ for each band, indicating that the instantaneous dynamical state carries substantial information about how strongly a node will respond. **(b)** Principal component analysis of feature-importance profiles across nodes. Left: loadings of the first principal components (PC1) for each band, revealing two regimes of time-specific mechanisms, one phase-gated, and another one based on the local synchrony and alignment. Right: cortical maps of PC1 scores for each band. **(c)** Effect of stimulation timing. For each band, we compare the response magnitude obtained when stimulating at randomly chosen time points versus at time points selected using the random forest predictions (“informed” stimulation). *** indicates p *<* 0.001

We next investigated whether frequency bands relied on similar time-specific features, and whether regions could be organised according to similar time-specific determinants of responsiveness. To this end, we performed a PCA across nodes on the feature-importance profiles. The first principal component (PC1) captured a dominant and highly consistent contrast across bands (**Figure 4b**). PC1 loaded strongly on the sine and cosine of the stimulated node’s local phase, whereas network-context features such as local synchrony and phase alignment loaded with the opposite sign (particularly in the faster bands). This defines two ends of the same axis, where nodes with high PC1 scores are primarily phase-gated, responding most when stimulated at specific phases, whereas nodes with low PC1 scores are more strongly modulated by network synchrony and alignment at the time of stimulation. Interestingly, slow rhythms (delta and theta) were predominantly phase-gated, whereas faster bands increasingly combined phase dependence with stronger modulation by network synchrony and alignment.

Finally, we tested whether these time-specific mechanisms can be used to optimise when to stimulate a given node. For each participant, node, and band, we applied the random forest to a set of candidate onset times along the baseline time series (held out from model training) to obtain a predicted response magnitude for each time point. We then compared two strategies, first, selecting an onset uniformly at random from this candidate set, and second, selecting the single top-ranked onset with the highest predicted response (“informed” stimulation). In all frequency bands, informed stimulation led to significantly larger responses than random stimulation (**Figure 4c**). This demonstrates that knowledge of the ongoing dynamical state can be used to systematically enhance the effectiveness of stimulation.

### 2.4 Dynamic node properties, more than target state, characterise the optimal nodes for the rest-to-listening transition

Once we had characterised each node’s response to external stimulation based on static and time-resolved features, we asked whether these same characteristics also determine which nodes are best to stimulate in order to achieve a brain-state transition, or whether optimal targets are instead primarily dictated by the target state itself.

For the rest-to-listening transition, we stimulated the resting state and quantified, for each node and band, how much this stimulation moved the resting FC pattern toward the listening-state FC. Specifically, the transition effect was the L2 distance between the stimulated node’s FC row (after stimulation in rest) and the FC row of the same node in the empirical listening state. For each participant and band, we selected the 15 nodes with the largest transition effect and counted how many of them belonged to an anatomically defined auditory network (see Methods), which we considered the network that would be expected a priori for this target state. As shown in **Figure 5a**, the number of optimal nodes falling in auditory cortex remained close to what would be expected by chance across all bands, indicating no systematic preference for auditory regions. The spatial distribution of optimal nodes across the cortex is displayed in **Figure 5b**, coloured by the percentage of participants for whom each node was selected as optimal.

**Figure 5:**
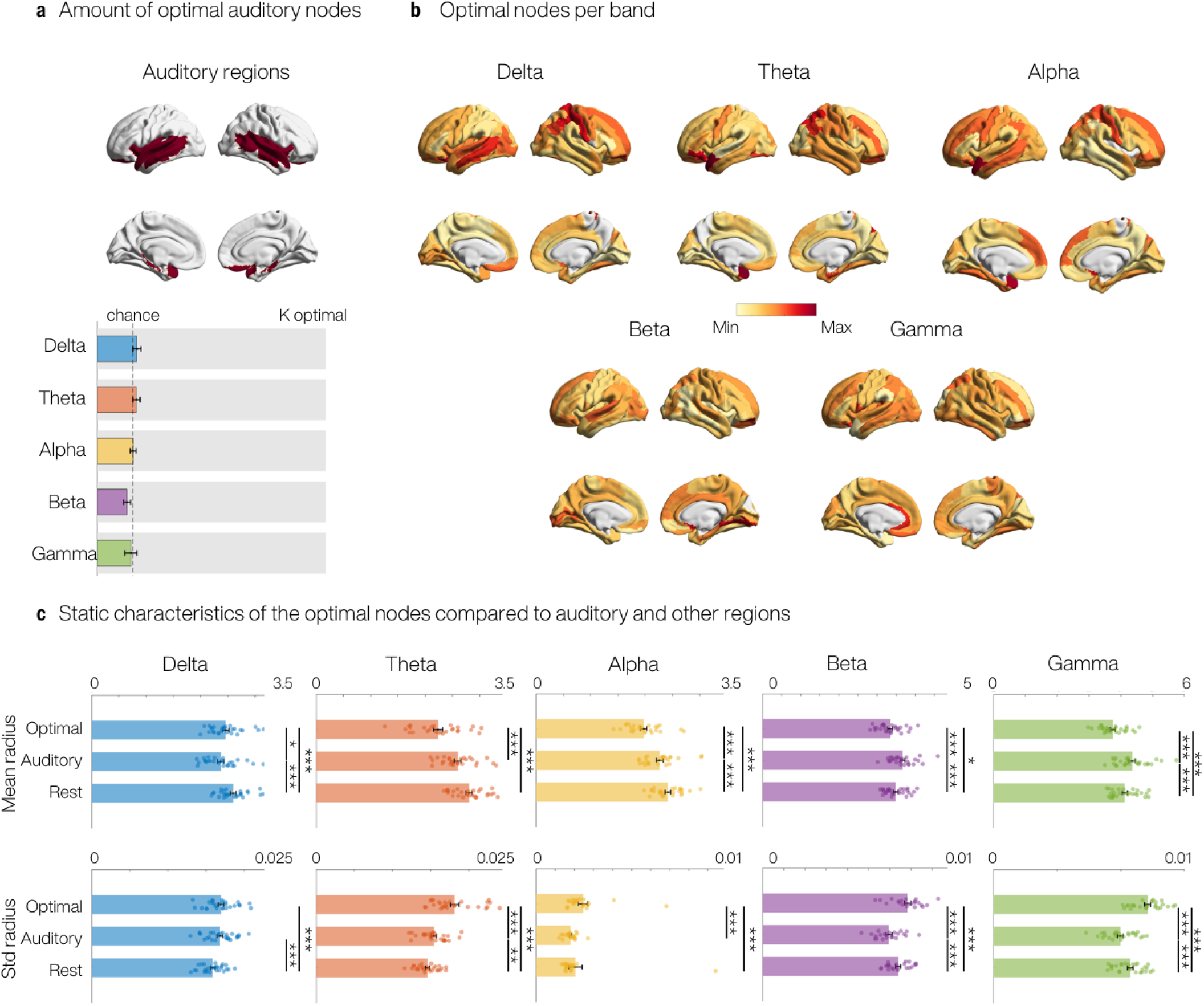
Optimal nodes for the rest-to-listening transition are defined by their baseline dynamics rather than by auditory membership. **(a)** Auditory network mask (top) and, for each band, number of optimal nodes (top 15 per participant) that fall within the auditory network. Coloured bars show the mean *±* SEM across participants; the grey line indicates chance level. **(b)** Cortical distribution of optimal nodes per band. Each node is coloured by the percentage of participants for whom it was selected among the 15 optimal nodes. **(c)** Baseline radius properties of optimal, auditory and remaining nodes. For each band, bars (mean *±* SEM) and dots (participants) show mean baseline radius (top row) and baseline radius standard deviation (bottom row) for optimal nodes, all auditory nodes and all remaining nodes. All p-values are FDR corrected, * indicates p *<* 0.05, ** indicates p *<* 0.01 and *** indicates p *<* 0.001

We then asked whether the baseline dynamical characteristics that previously explained response magnitude (mean baseline radius and radius variability) also distinguished the optimal nodes from the rest of the cortex, and whether this differed for auditory versus non-auditory regions. Recall that smaller mean radius and larger radius variability were associated with stronger responses to stimulation. As shown in **Figure 5c**, across all frequency bands optimal nodes had a significantly lower mean radius and a significantly higher radius standard deviation than both auditory nodes and the remaining non-auditory nodes (all p-values are FDR corrected; * indicates p *<* 0.05, ** indicates p *<* 0.01 and *** indicates p *<* 0.001). Overall, this indicates that the nodes that best drive the rest-to-listening transition are not defined primarily by belonging to the target (auditory) network, but rather by a dynamical regime characterised by low baseline radius and high radius variability.

## 3 Discussion

In this study, we asked which static (time-aggregated summaries that are constant across stimulation onsets, e.g., structural connectivity strength) and time-resolved (features computed at the exact onset, e.g., phase) properties determine how strongly a region responds to stimulation. We also tested whether the regions that most effectively drive a transition between brain states (rest to listening) are task-relevant (e.g. auditory cortex) or instead determined by more general dynamical mechanisms. Using band-specific Hopf whole-brain models fitted to MEG data, we found that nodes with lower baseline phase-space radius and higher temporal variability of this radius showed the largest responses to stimulation. We further found that responsiveness is time-dependent, with some regions depending primarily on instantaneous local phase at the time of stimulation, whereas others are more strongly modulated by network synchrony and alignment. Crucially, when targeting the rest-to-listening transition, the most effective stimulation sites are not consistently restricted to auditory cortex, but are instead characterised by the same low-radius and high-variability properties that predict responsiveness. Together, these results suggest that where and when to stimulate is shaped by state-, band-, and dynamics-dependent features.

The first step was to build a whole-brain model for each participant and state (rest or listening). Specifically, we used a Hopf model [26] and fitted it separately to each canonical frequency band of the MEG data. To capture directed interactions between brain areas beyond the structural connectome alone, we optimised the model to estimate a generative effective connectivity (GEC) matrix for each participant, state, and band. Previous studies have also used whole-brain models to simulate MEG data, but typically either relied on the structural connectome [27, 28, 29, 30, 31, 32], or fitted a single effective connectivity across the full spectrum without band-specific estimation [33]. Although some previous studies have modelled individual frequency bands separately [29, 27, 30], we extend this line of work by estimating band-specific GEC, which, to our knowledge, has not been done previously. This band-wise approach allows us to isolate how the system responds within distinct oscillatory regimes, which is essential given that brain networks operate differently across frequencies [34]. Moreover, optimising one GEC for each band enhances the model’s ability to reproduce the empirical data in each band, going beyond the structural connectome by capturing directed effective interactions.

Having accurately simulated the empirical data, we used the model to apply in silico perturbations and examine how each frequency band responded (as quantified by stimulation-induced changes in functional connectivity). We found that the delta band elicited the strongest responses in the resting state, whereas the alpha band dominated during the listening condition. The heightened delta response in the resting state likely reflects the dominance of slow oscillatory modes in the absence of external demands [35, 36]. Delta rhythms are thought to organize large-scale spontaneous fluctuations and resting-state functional connectivity, naturally predisposing the system to respond more strongly to perturbations in this range. This aligns with evidence that stimulating near the brain’s endogenous frequency enhances its responsiveness [37]. In contrast, the increased alpha-band stimulation–evoked FC reconfiguration during the listening task may relate to task-dependent modulation of alpha rhythms, including alpha-band event-related desynchronization (ERD). When the brain is engaged in a task, relevant regions exhibit a reduction in alpha power, which has been linked to increased excitability [38, 39, 40]. Thus, stimulation in the alpha band might interact with an already highly excitable state, amplifying the brain’s responsiveness compared to frequency bands that are not actively modulated by the task. Supporting this, in vivo studies have shown that alpha-band stimulation during tasks can modulate ERD and improve behavioral performance [41, 42]. Together, these results suggest that task engagement can potentiate alpha-band stimulation effects by aligning stimulation with an intrinsically responsive dynamical regime [43].

We also found that the regions that respond most strongly to stimulation are those with low baseline radius and high radius variability. Here, radius refers to the phase-space radial amplitude of the Hopf state, 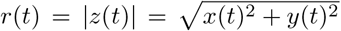, which quantifies the instantaneous oscillation amplitude of each node. Low baseline radius corresponds to trajectories concentrated near the origin in the Hopf phase space [44, 45]. In the Hopf model, the amplitude is stabilised by a nonlinear saturation term (often written as 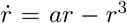), meaning that when baseline *r* is small the amplitude-dependent restoring effect is weaker and external inputs, including stimulation, can induce larger deviations [46, 47, 45]. In addition, radius variability was associated with higher responsiveness. Nodes whose radius fluctuates more over time visit low-amplitude regimes more frequently, increasing the likelihood of time windows in which perturbations elicit particularly large responses. This amplitude-based account complements previous work linking transition-like regimes to heightened susceptibility and prolonged recovery following perturbations (critical slowing down) [48, 49, 44, 50, 51, 52]. Finally, although structural features such as eigenvector centrality or degree did not directly predict response strength, they may still shape a region’s dynamical context by modulating recurrent input and effective drive, thereby indirectly influencing baseline radius and its variability [45, 21].

Another interesting result was that the resting state consistently showed stronger correlations between responsiveness and the amplitude features (baseline radius and radius variability) than the listening state. This likely reflects the fact that rest is more strongly governed by spontaneous fluctuations, allowing local dynamical properties to exert a greater influence on how regions respond to stimulation [44, 29, 53]. In contrast, task engagement can reduce intrinsic dynamical variability [54] and alter regional excitability [55], potentially attenuating the impact of these baseline features on perturbation responses.

Once we understood the mechanisms explaining which regions respond more to stimulation, we turned to time-resolved features that predict when regions respond most strongly. Using random-forest models trained on instantaneous local features, we found that time-varying responsiveness can be organised into two dominant regimes. In a phase-driven regime, response magnitude is primarily gated by the stimulated node’s instantaneous local phase at the time of stimulation. In a network-driven regime, response magnitude is more strongly modulated by network context, including synchrony and phase alignment with surrounding activity. These regimes were not equally prevalent across frequency bands: slower bands (delta and theta) were predominantly phase-driven, whereas faster bands showed an increasing contribution of network-driven factors. The idea that the brain’s instantaneous state modulates stimulation efficacy is gaining support [25, 10], and our results align with empirical work showing that stimulation responses depend not only on local phase but also on inter-regional phase coupling [56, 57, 58, 59]. Consistent with these mechanisms, selecting stimulation times based on the ongoing dynamics reliably increased response magnitude relative to random timing. Altogether, these findings support the potential of closed-loop or informed stimulation protocols [60].

Finally, we asked whether the regions best suited to drive a transition from rest to a listening state were defined primarily by their anatomical role in the target network (auditory regions), or by intrinsic dynamical properties. We found that optimal stimulation targets were not consistently located in auditory cortex, instead, across frequency bands, the most effective nodes had low baseline radius and high radius variability, the features that predicted strong responsiveness to perturbation. In other words, a node’s ability to drive the model toward a target state depends less on its predefined functional label and more on whether it operates in a dynamical regime that enables large stimulation-induced reconfiguration. This perspective is broadly consistent with prior work suggesting that stimulation of specific control-relevant nodes can be effective for driving state changes, sometimes independently of the cognitive domain of the target state [61, 62]. Complementarily, empirical and clinical studies have shown that targeting functionally relevant regions can improve performance in visual, auditory, or somatosensory tasks [63, 64, 65]. However, those targets are typically chosen a priori based on presumed functional relevance, and although relevant for the transition, they might not be the most effective ones. Our results highlight that model-based analyses can move beyond a priori assumptions by providing a mechanistic route to identifying stimulation targets, which may differ from task-local regions and yield new candidates for inducing desired brain-state transitions.

A natural question is why stimulating non-auditory nodes can nonetheless reproduce a rest-to-listening transition, and whether such stimulation would instead drive the brain toward other sensory task states. In our framework, “listening” is defined operationally as the empirically observed listening-state functional connectivity, and stimulation targets were selected to maximize similarity to this specific pattern (not to induce a generic task-like change). That the most effective stimulation sites are not consistently located within auditory cortex therefore does not imply that auditory regions are irrelevant. Instead, in a coupled network, the most effective place to stimulate to steer the whole system toward a target pattern need not coincide with the regions that most prominently characterise that pattern [66]. Moreover, stimulation effects can spread beyond the stimulated site and induce distributed network reconfiguration [67, 68], which provides a plausible route by which stimulation outside auditory cortex can still shift whole-brain connectivity toward the listening pattern. At the same time, we did not test transitions to other task-defined targets (e.g., visual or somatosensory states), so we cannot claim that the identified stimulation sites are uniquely listening-specific, which is an important direction for future work.

There are several limitations that should be acknowledged. First, our conclusions are derived from a computational model that necessarily simplifies brain dynamics. While this supports mechanistic interpretation, the Hopf model provides a simplified description of whole-brain oscillatory dynamics that is fitted to reproduce empirical functional and time-shifted covariance, capturing key band-limited structure and large-scale coupling. However, it does not uniquely identify the underlying biological mechanisms and necessarily abstracts away microcircuit processes (e.g., spiking and synaptic dynamics) as well as richer whole-brain features not fixed by second-order statistics (e.g., cross-frequency interactions and nonstationary switching), which may also influence how stimulation effects emerge and propagate. Second, although we find robust associations between stimulation responsiveness and radius-related quantities (baseline radius and its variability), we do not demonstrate that manipulating radius alone is sufficient to reproduce the observed effects, since parameters can covary and stimulation responses may depend on interactions between local dynamics and coupling. Third, the model’s network topology (effective connectivity) is treated as fixed, meaning that we did not test whether alternative connectomes, or systematic perturbations of topology, would preserve the same qualitative conclusions. This leaves open the question of how brain-specific versus model-specific these effects are, and whether steerability patterns would change under different structural constraints. Fourth, the simulated perturbations are simplified and do not account for key biophysical constraints (e.g., field spread, neuronal orientation, heterogeneous tissue conductivity), which limits direct clinical actionability at present. Fifth, responsiveness was quantified via stimulation-induced changes in functional connectivity centred on the stimulated node, which captures how stimulation reshapes a node’s interaction profile, but does not fully characterise whole-network reorganisation. Sixth, fitting separate band-specific models isolates oscillatory regimes but omits cross-frequency interactions that may influence stimulation responses. Finally, the model is built from MEG source-reconstructed data, which has known limitations including limited spatial resolution, sign ambiguity in source reconstruction, and reduced sensitivity to deep or radially oriented sources [69, 70].

These limitations motivate several methodological and experimental extensions. On the modelling side, targeted sensitivity analyses could test causality by manipulating radius-related parameters independently of coupling and other covarying quantities, and by assessing robustness across alternative structural or effective connectomes. Incorporating more realistic stimulation models (e.g., forward fields, spatial spread, and tissue heterogeneity) and expanding outcome measures beyond node-centred FC changes to whole-network reconfiguration would help bridge the gap to in vivo settings. Experimentally, the predictions could be evaluated against datasets with known stimulation timing and targets (e.g., TMS–EEG/MEG, intracranial stimulation, or DBS recordings), with particular emphasis on whether state- and frequency-dependent “steerable regimes” generalise across individuals and tasks.

Overall, this study characterises the spatial and temporal control landscape of stimulation in personalised whole-brain models. We show that responsiveness is strongest in nodes with low baseline radius and high radius variability, that stimulation is strongly timing-dependent through phase- and network-gated regimes across frequency bands, and that targets that most effectively drive rest-to-listening transitions are better predicted by dynamical response properties than by task anatomy alone. Together, these findings suggest that stimulation efficacy may be usefully conceptualised as a network-dynamics problem, shifting emphasis from asking where the function is localised to asking which network-level dynamical regimes are most steerable, and when.

## 4 Methods

### 4.1 Participants

The study included 29 participants (17 female, 12 male), aged 19–50 years (mean age = 26.45 ± 7.37 years). Four participants were first-year music students, and the remaining participants were non-musicians. All participants, except one non-musician, had *<* 2 years of formal musical theory training (mean years = 0.23 ± 0.47) and *<* 5 years of formal musical instrument training (mean years = 0.59 ± 1.10). None of the participants reported any neurological or psychiatric disorder, and none reported hearing impairment. Only one participant was left-handed. Participants came from 14 different countries, with Danish participants forming the largest group (24%). Participants were compensated with online shopping vouchers. Two participants did not complete the passive listening recording, and one additional participant was excluded due to technical problems during data acquisition, resulting in a final sample of N = 26 with complete data in both conditions.

### 4.2 Experimental procedure

Participants completed MEG recordings in two experimental conditions. During passive listening, they heard a rhythmic auditory sequence while watching a silent movie. The sound sequence comprised repeated 300 Hz sine tones (100 ms each, with 10 ms onset/offset ramps) delivered at 2.4 Hz (inter-onset interval 410 ms) for 5 minutes. Before recording, stimulus intensity was individually calibrated to 50 dB above each participant’s hearing threshold. During rest, participants watched the same silent movie without auditory stimulation for 10 minutes; for comparability with the passive listening condition, analyses were restricted to the first 5 minutes of the resting-state recording.

The data analysed here were drawn from a larger dataset collected at Aarhus University. The study received approval from the Ethics Committee of the Central Denmark Region (De Videnskabsetiske Komi-teer for Region Midtjylland; ref. 1-10-72-411-17) and was conducted in accordance with the Declaration of Helsinki.

### 4.3 Data acquisition

MEG recordings were acquired at Aarhus University Hospital in Aarhus, Denmark, inside a magnetically shielded room, using an Elekta Neuromag TRIUX system with 306 sensors (Elekta Neuromag, Helsinki, Finland). Signals were sampled at 1000 Hz and recorded with online analog filtering between 0.1–330 Hz. Before the session, each participant’s head shape was digitized and the locations of four head-position indicator (HPI) coils were measured relative to three anatomical landmarks using a 3D digitizer (Polhemus Fastrak, Colchester, VT, USA). These digitization data were later used for MEG–MRI co-registration. Throughout acquisition, HPI coils provided continuous head-position tracking to support motion correction during preprocessing. In addition, bipolar electrocardiography (ECG) and electrooculography (EOG) signals were recorded to enable identification and removal of cardiac- and ocular-related artifacts.

Structural MRI data were collected on a separate day using a CE-approved 3T Siemens scanner at the same site. High-resolution T1-weighted anatomical images (MPRAGE with fat saturation) were acquired at 1.0 × 1.0 × 1.0 mm resolution, with the following parameters: TE = 2.61 ms, TR = 2300 ms, reconstructed matrix 256 × 256, echo spacing 7.6 ms, and bandwidth 290 Hz/Px.

### 4.4 MEG data pre-processing

Raw MEG data (204 planar gradiometers, 102 magnetometers) were first cleaned using MaxFilter [71] to attenuate external interference via spatiotemporal signal space separation (tSSS). Data were down-sampled from 1000 Hz to 150 Hz and corrected for head motion using continuous HPI-based movement compensation (default 10 ms step). A correlation limit of 0.98 between inner and outer subspaces was used within tSSS to reject intersecting signals. Subsequent preprocessing was performed in MATLAB/SPM format using the Oxford Centre for Human Brain Activity (OHBA) Software Library (OSL) [72] and custom scripts (LBPD, https://github.com/leonardob92/LBPD-1.0.git). Continuous recordings were visually inspected to remove large artifacts, and independent component analysis was applied to suppress ocular and cardiac artifacts: components were identified based on elevated correlation with EOG/ECG channels and confirmed by inspection of their time courses and sensor topographies before removal. Cleaned sensor data were reconstructed by back-projecting the remaining components. For each condition, data were analysed as a single 5-minute segment. For full implementation details, see [73].

### 4.5 Source reconstruction

Sensor-level MEG data were projected to source space using a beamforming approach implemented with routines from OSL/SPM/FieldTrip together with in-house MATLAB code [74, 75, 76]. Source reconstruction followed the standard forward–inverse framework. For the forward model, individual T1-weighted anatomical scans were co-registered to MEG space using fiducial landmarks and digitized head-shape points; when an individual MRI was unavailable, a standard MNI152 template (8 mm resolution) was used instead. A single-shell head model (Nolte method) was used to compute the lead field [77].

Source space was discretized on an 8 mm grid (3559 locations), and reconstruction was performed using magnetometer channels only, motivated by their higher sensitivity to deeper sources relative to planar gradiometers [78]. For the inverse solution, we used a linearly constrained minimum variance (LCMV) beamformer. Beamformer weights were derived from the sensor covariance estimated on the continuous recording, separately for each experimental condition. Dipole orientations were initially modelled in three orthogonal directions and then reduced to a single dominant orientation per location using an singular value decomposition (SVD)-based projection prior to weight computation. To reduce depth-related bias in reconstructed power, beamformer weights were normalized [79]. The normalized weights were finally applied to the neural activity, resulting in voxel-wise source time series obtained for the full analysed segment for each condition [74]. For full implementation details and parameter settings, see [73].

### 4.6 Data parcellation

Source-level activity was parcellated using the Automated Anatomical Labeling (AAL) atlas. AAL labels were mapped onto the 8-mm source grid in MNI space, and for each AAL parcel we identified the set of source-grid voxels assigned to that parcel. For each parcel, we extracted all voxel time series and computed the pairwise Pearson correlation matrix across voxels. We then selected the voxel with the largest sum of absolute correlations to all other voxels in the same parcel (excluding the diagonal) and used its time series as the parcel representative. Parcellation resulted in *N* = 90 regional time series (AAL atlas).

For stratified analyses that distinguished auditory-related regions, we defined an a priori set based on AAL labels including Heschl’s gyrus and superior temporal gyrus parcels. In addition, we grouped other unimodal sensory–motor regions using the same label-based approach, including visual-related cortex (Occipital, Calcarine, Cuneus, Lingual, Fusiform) and somatomotor-related cortex (Precentral, Postcentral, Rolandic Operculum).

### 4.7 Whole-brain modelling

#### 4.7.1 Band-specific time series

All whole-brain modeling analyses were performed independently within the five canonical frequency bands (Delta: 1–4 Hz; Theta: 4–8 Hz; Alpha: 8–12 Hz; Beta: 13–30 Hz; Gamma: 30–80 Hz). For each subject and condition, we divided the parcellated time series into band-specific signals by band-pass filtering within these ranges.

#### 4.7.2 Model definition

To simulate the activity of individual brain areas, we employed the Stuart–Landau oscillator model, which characterizes each region by dynamics near a supercritical Hopf bifurcation. This bifurcation enables transitions from noisy fluctuations to oscillatory behaviour [80]. Models based on this principle have effectively reproduced critical features of neural dynamics across modalities such as electroencephalography [81, 82], magnetoencephalography [83], and functional magnetic ressonance imaging [84, 85].

In this approach, for each frequency band, the brain is modeled as a network of *N* coupled oscillators, each governed by the Stuart–Landau equation and interacting through the connectivity matrix *C*:

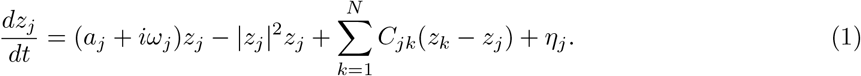

Here, *z*_*j*_ = *x*_*j*_ + *iy*_*j*_ denotes the complex state of region *j, η*_*j*_ is Gaussian noise with variance *σ*^2^, and *a*_*j*_ is the bifurcation parameter. When *a*_*j*_ *<* 0, the system stabilizes at a fixed point at *z* = 0, whereas *a*_*j*_ *>* 0 leads to self-sustained oscillations with frequency 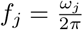.

Since the model requires specifying an intrinsic frequency for each node, we estimated *ω*_*j*_ empirically from the band-limited MEG time series. Specifically, for each frequency band, we computed the power spectrum of each region (after *z*-scoring the time series) and identified the region-specific peak frequency within the corresponding band. This yielded *f*_*j*_ per region, and we set *ω*_*j*_ = 2*πf*_*j*_.

#### 4.7.3 Model linearization

Following earlier work [86], we selected *a* = −0.02 to position the system close to the bifurcation point, facilitating a linear approximation suitable for analytical treatment of functional connectivity [45]. Under this linear noise approximation (LNA), the system dynamics in vectorized form are described as:

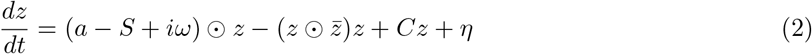

where **z** = [*z*_1_, …, *z*_*N*_]^*T*^, **a** = [*a*_1_, …, *a*_*N*_]^*T*^, ***ω*** = [*ω*_1_, …, *ω*_*N*_]^*T*^, ***η*** = [*η*_1_, …, *η*_*N*_]^*T*^, and **S** = [*S*_1_, …, *S*_*N*_]^*T*^, where **S** is a vector indicating the strength of each node, with *S*_*i*_ = ∑_*j*_ *C*_*ij*_. The transpose operation is denoted by []^*T*^, the ⊙ symbol represents the Hadamard element-wise product, and 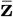 is the complex conjugate of **z**.

Linearizing around the stable solution **z** = 0, we obtain the system for small perturbations:

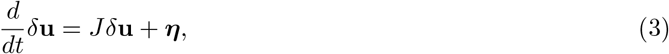

where the 2*N* -dimensional vector *δ***u** = [*δ***x**, *δ***y**]^*T*^ = [*δx*_1_, …, *δx*_*N*_, *δy*_1_, …, *δy*_*N*_]^*T*^ includes the fluctuations in both the real and imaginary parts of the state variables. The 2*N* × 2*N* matrix *J* stands for the Jacobian of the system computed at the steady state:

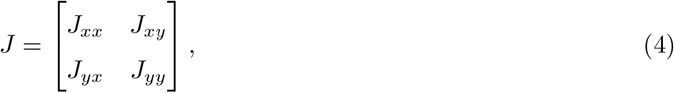

with *J*_*xx*_, *J*_*xy*_, *J*_*yx*_, and *J*_*yy*_ being matrices of size *N* × *N* . Specifically, *J*_*xx*_ = *J*_*yy*_ = diag(**a** − **S**) + *C* and *J*_*xy*_ = −*J*_*yx*_ = diag(***ω***). The validity of this approximation relies on the stability of the origin.

To derive the covariance matrix *K* = ⟨*δ**uδu***^*T*^⟩, we reformulate Equation 3 as *dδ***u** = *Jδ***u** *dt* + *d***W**, where *d***W** is a 2*N* -dimensional Wiener process with covariance ⟨*d**WdW***^*T*^⟩ = *Q dt*. Here, *Q* is the covariance matrix of the noise. Using Itô’s stochastic calculus, we obtain:

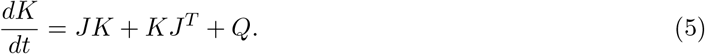

This allows us to compute the stationary covariances (for which 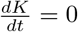) by

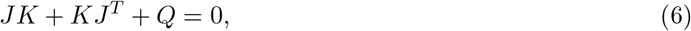

which is a Lyapunov equation that can be solved through the eigen-decomposition of the Jacobian matrix *J* [45]. By extracting the first *N* rows and columns of *K* and taking the real part, we obtain the simulated zero-lag covariance of the real components,

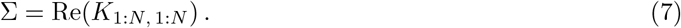

The simulated functional connectivity is then computed as the normalized covariance (Pearson correlation),

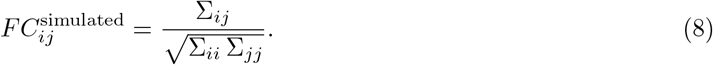

In addition, simulated time-shifted covariances were obtained analytically as

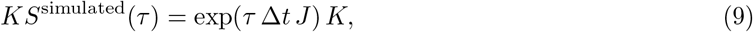

from which we extracted 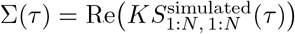 and normalized analogously to yield

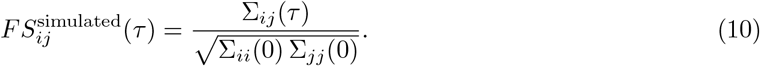

The time lag was chosen as a short delay in the band-limited signals, using *τ* = 3 samples for delta– alpha and *τ* = 2 samples for beta and gamma, guided by the faster autocorrelation decay observed at higher frequencies.

#### 4.7.4 Optimization of the model to compute GEC

To match the model to empirical band-limited MEG data, we refined the connectivity matrix *C* through an iterative pseudo-gradient optimization method. The resulting optimized matrix, named as the generative effective connectivity (GEC), reflects directional influences between regions more accurately than structural connectivity derived from diffusion MRI. The optimization aimed to reduce the discrepancy between simulated (FC^simulated^) and empirical functional connectivity (FC^empirical^), as well as the empirical and simulated time-shifted covariances (FS^empirical^(*τ*), FS^simulated^(*τ*)), where *τ* indicates the time lag. These were computed by shifting the empirical covariance matrix (KS^empirical^(*τ*)) and normalizing each pair (*i, j*) by 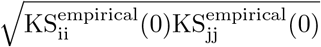. Notably, incorporating time-shifted covariances can result in asymmetries within the coupling matrix *C*.

The update rule for *C* at each iteration was:

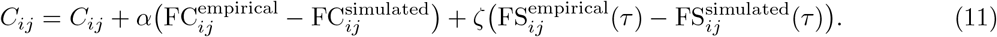

The fitting of the model is iteratively executed until the updated value of *C* reaches a stable value, indicating that the model is optimized. The initial value of *C* is set using the anatomical connectivity dMRI data obtained through probabilistic tractography, and it is only updated if connections exist in this matrix. This procedure is followed rigorously, except for homologous connections between regions in the two hemispheres, which are also updated, as tractography is less reliable for this type of connectivity. Additionally, *α* and *ζ* are set to 0.001. The final matrix, GEC, characterizes the brain’s effective connectivity under the modeled dynamics [87].

### 4.8 In silico stimulation protocol

#### 4.8.1 Baseline nonlinear simulations from fitted GEC

For each participant, frequency band, and condition, we first estimated a generative effective connectivity matrix (GEC) using the linear noise approximation of the Hopf/Stuart–Landau model explained above. We then used the resulting GEC as the coupling matrix in the nonlinear Hopf model to simulate baseline timeseries.

Each brain region *j* was represented by a two-dimensional state (*x*_*j*_(*t*), *y*_*j*_(*t*)) governed by the nonlinear Hopf dynamics with diffusive coupling:

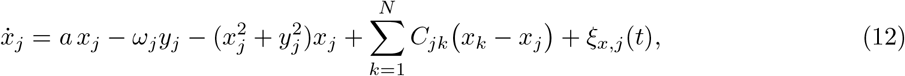

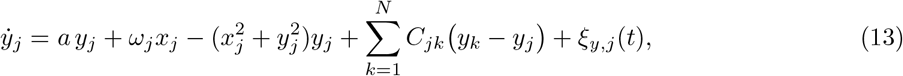

where *a* = −0.02 for all nodes, *C* denotes the participant/band/condition-specific GEC, and *ω*_*j*_ = 2*πf*_*j*_ uses the region-specific intrinsic frequency estimated from the empirical band-limited signals (same as before). Independent Gaussian noise was added to both components with amplitude *σ* = 0.01.

Simulations were performed at an output sampling rate of *f*_*s*_ = 150 Hz (Δ*t* = 1*/f*_*s*_), using an Euler– Maruyama integration step *dt* = 0.1 Δ*t/*2. For each simulation, we first ran a burn-in period of 1000 s to remove transients. We then simulated *T*_max_ = 9000 samples (60 s at 150 Hz) and stored both the baseline time series *x*_*j*_(*t*) and the full phase-space trajectories (*x*_*j*_(*t*), *y*_*j*_(*t*)) for each region. These baseline trajectories were subsequently used to define stimulation onset states and to compute the baseline functional connectivity reference.

#### 4.8.2 Sinusoidal perturbations

To quantify how stimulation effects depend on the ongoing brain state, we applied node-wise sinusoidal perturbations at multiple onset times extracted from the baseline trajectory. For each participant, band, and node *i*, we selected onset indices with a fixed stride of 500 samples (i.e., onsets every 500 Δ*t* along the baseline trajectory). At each onset *t*_0_, we initialized a new simulation from the saved baseline phase-space state **z**(*t*_0_) = [*x*(*t*_0_), *y*(*t*_0_)] and applied an external sinusoidal drive to the node *i*:

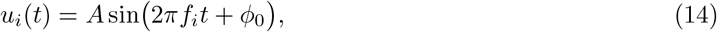

where *f*_*i*_ is the intrinsic frequency of node *i* in the corresponding band, and the initial drive phase was fixed to *ϕ*_0_ = 0 for all onsets. The stimulation amplitude was explored over a grid *A* ∈ {10, 20, 30, 40, 50, 60, 70}, and as results did not vary, finally kept A = 10 for the analyses.

For each onset, we simulated a post-onset segment of *T*_max_ = 2000 samples, with stimulation applied for the first 1000 samples and then switched off for the remaining samples.

#### 4.8.3 Functional connectivity and effect metrics

For each participant, band, and condition, we computed a baseline functional connectivity matrix *FC*^base^ from the baseline simulated time series (Pearson correlation across regions; diagonal set to zero). For each stimulated simulation (node *i*, onset *t*_0_), we computed the corresponding connectivity matrix *FC*^stim^(*i, t*_0_) and quantified stimulation-induced changes relative to baseline.

Our primary node-level effect metric was the row-wise *L*_2_ change in connectivity of the stimulated node:

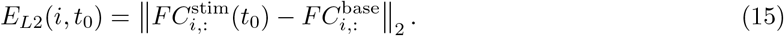

#### 4.8.4 Static node metrics

To relate stimulation effects to time-invariant node properties, we computed a set of metrics for each participant, frequency band, and condition. Static network metrics were derived from the fitted GEC, denoted *GEC* ∈ *R*^*N×N*^, and dynamical metrics were derived from the baseline nonlinear Hopf simulations (phase-space trajectories).

##### Weighted in- and out-strength

Given the directed weighted matrix *GEC*, we defined the in-strength of node *i* as the sum over the *i*-th row,

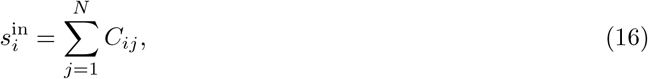

and the out-strength as the sum over the *i*-th column,

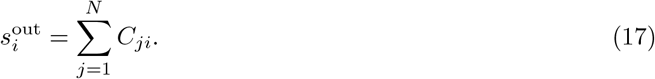

##### Eigenvector centrality

Eigenvector centrality was defined as the dominant eigenvector of *GEC* (largest eigenvalue), and we took its absolute value for stability across sign conventions.

##### Mean and variability of radius

From the baseline nonlinear Hopf simulations we obtained phase-space trajectories (*x*_*i*_(*t*), *y*_*i*_(*t*)) for each node. We defined the instantaneous radius

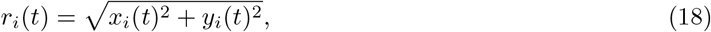

and computed its temporal mean and standard deviation:

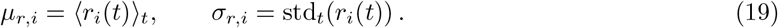

#### 4.8.5 Time-resolved stimulation-state metrics

To characterize the dynamical state at the moment of stimulation, we computed a set of time-resolved features at each stimulation onset *t*_0_ from the baseline phase-space state. Let *z*_*j*_(*t*) = *x*_*j*_(*t*) + *i y*_*j*_(*t*) denote the complex state at time *t*, with phase *ϕ*_*j*_(*t*) = atan2(*y*_*j*_(*t*), *x*_*j*_(*t*)) and radius *r*_*j*_(*t*) = |*z*_*j*_(*t*)|.

For each target node *i* and onset *t*_0_, we computed the following 12 features:

##### Local phase and amplitude

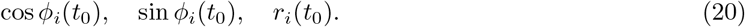

##### Instantaneous coupling input

We defined the complex coupling input to node *i* as

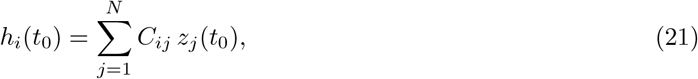

and extracted its magnitude |*h*_*i*_(*t*_0_)|. We also computed the radial component of this input, i.e., the projection of *h*_*i*_(*t*_0_) onto the direction of *z*_*i*_(*t*_0_):

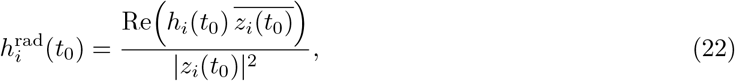

with 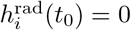 when |*z*_*i*_(*t*_0_)| = 0.

##### Global synchrony and mean radius

We computed a Kuramoto-like global order parameter from the instantaneous phases:

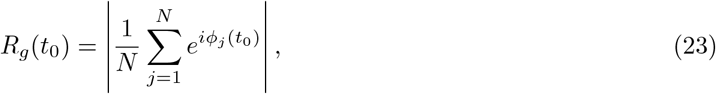

and the mean global radius 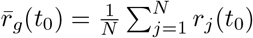. We also quantified alignment of the stimulated node with the global phase as

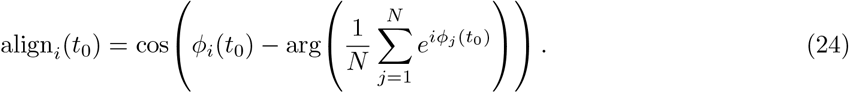

##### Local network synchrony and radius

Using the outgoing weights of node *i, w*_*ij*_ = *C*_*ij*_, we formed normalized weights 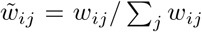 when ∑_*j*_ *w*_*ij*_ *≠* 0 (and set local features to 0 otherwise).

We then computed a local phase coherence

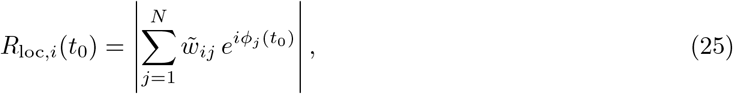

and a local weighted mean radius

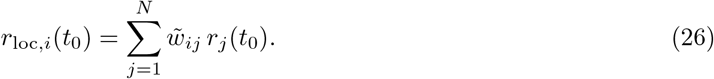

##### History features

Finally, we computed two history terms over a pre-onset window of length *L* samples (here *L* = 45) immediately preceding *t*_0_:

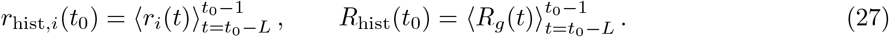

#### 4.8.6 Random-forest modelling of time-resolved predictors

To quantify how time-resolved stimulation-state features predict onset-by-onset stimulation effects, we trained a random-forest regressor separately for each participant, frequency band, and stimulated node. Each training sample corresponded to a stimulation onset *t*_0_ and consisted of the 12 time-resolved predictors evaluated at *t*_0_, with the response given by the stimulation effect *E*(*i, t*_0_) (row-wise *L*_2_ FC change). Random forests were implemented in MATLAB (TreeBagger) using 300 trees (regression mode) and out-of-bag (OOB) prediction. Predictive performance was quantified using OOB predictions to compute the coefficient of determination (*R*^2^). Feature relevance was quantified using permutation-based OOB predictor importance (OOBPermutedPredictorDeltaError), yielding one importance value per feature aggregated across onsets for each node.

### 4.9 Statistical analysis

All statistical analyses were performed in MATLAB using two-sided non-parametric tests with *α* = 0.05. Differences between paired measurements (e.g., rest vs listening and between frequency bands within subject) were assessed using Wilcoxon signed-rank tests (signrank). Associations between node stimulation effects and node metrics were quantified using Spearman rank correlation across nodes within each subject. Multiple comparisons were controlled using the Benjamini–Hochberg false discovery rate (FDR) procedure, applied separately within each condition for condition-specific analyses. For exploratory dimensionality reduction of the 12-dimensional random-forest importance profiles, we performed PCA separately within each frequency band after z-scoring features across samples.

## 5 Data and Code Availability

The data is available in the following Zenodo repository: https://doi.org/10.5281/zenodo.14917255. The code that supports the findings of this study are available from the corresponding author upon request.

## 6 Authors Contributions

I.A.-P. conceived the study, developed the methodology, implemented the software, performed the formal analysis and investigation, prepared the visualizations, and wrote the original draft of the manuscript, with further review and editing. L.B. contributed to conceptualization and methodology, provided resources, supervised the work, and reviewed and edited the manuscript. M.R. contributed to conceptualization, provided resources, and reviewed and edited the manuscript. Y.S.-P. reviewed and edited the manuscript. P.V. and M.L.K. provided resources and reviewed and edited the manuscript. G.D. contributed to methodology, supervised the work, acquired funding, and reviewed and edited the manuscript.

## 7 Declaration of Competing Interests

The authors declare no competing interests.

## 8 Aknowledgements

I.A.-P. was supported by grant PID2022-136216NB-100 funded by MICIU/AEI/10.13039/501100011033 and by the European Regional Development Fund (ERDF), “A way of making Europe” (EU). L.B. was supported by the Sapere Aude: Independent Research Fund Denmark (DFF) Research Leader grant (grant ID: 10.46540/5253-00003B), the Center for Music in the Brain, and Linacre College, University of Oxford. M.R. was supported by the Center for Music in the Brain and the Nordic Mensa Fund. Y.S.-P. was supported by the ERC Synergy grant NEurological MEchanismS of Injury, and Sleep-like cellular dynamics (NEMESIS; ref. 101071900), funded by the European Union (Horizon Europe). M.L.K. was supported by the Centre for Eudaimonia and Human Flourishing, funded by the Pettit and Carlsberg Foundations, and by the Center for Music in the Brain. G.D. was supported by grant PID2022-136216NB-I00 funded by MICIU/AEI/10.13039/501100011033 and by the European Regional Development Fund (ERDF), “A way of making Europe” (EU); by the ERC Synergy grant NEMESIS (ref. 101071900), funded by the European Union (Horizon Europe); and by the AGAUR research support grant 2021 SGR 00917 funded by the Department of Research and Universities of the Generalitat of Catalunya. The Center for Music in the Brain is funded by the Danish National Research Foundation (project number DNRF117), the Lundbeck Foundation (R469-2024-1573), and Købmand Herman Sallings Fond. We thank Simjon Radloff, Anastasiia Popova, and Antonieta Martínez-Guerrero for their extensive and tireless work during data collection

## A Supplementary Material

**Figure S1:**
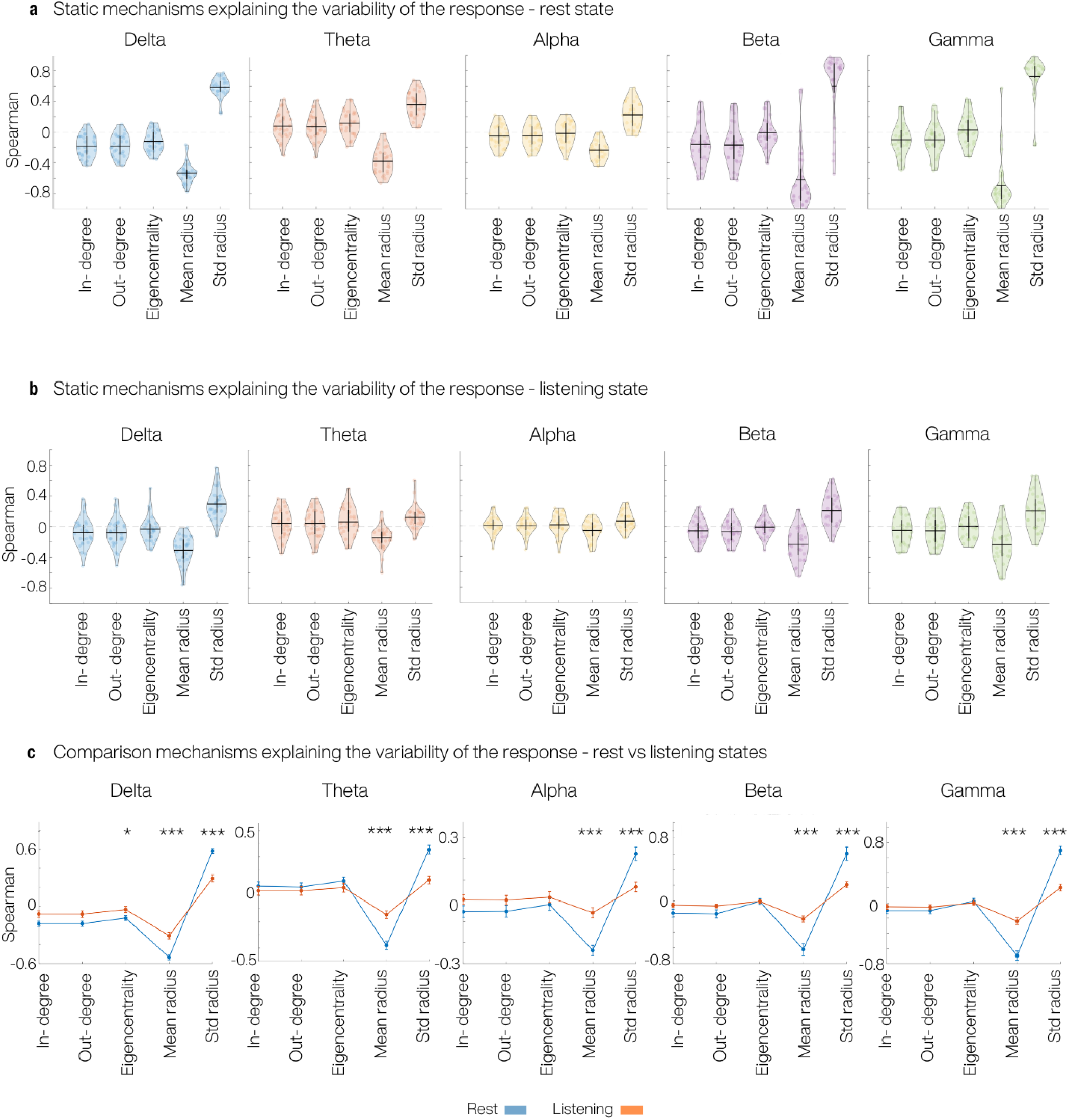
Node dynamics explains response variability over time after stimulation. **(a)** Distribution across participants of Spearman correlations between node-wise response magnitude variability and five static metrics (in-degree, out-degree, eigenvector centrality, mean baseline radius and baseline radius variability) for each frequency band in the resting state. **(b)** Same as in (a), but for listening state instead of the resting state. **(c)** Direct comparison of mechanisms between rest and listening for the response magnitude variability. Lines show the mean *±* SEM of the Spearman correlations across participants, separately for each band and metric, asterisks indicate significant rest–listening differences (FDR-corrected, * indicates p *<* 0.05 and *** indicates p *<* 0.001). Across bands, mean radius correlates negatively and std radius positively with the magnitude of the response, with systematically stronger effects at rest than during listening.

## Notes

### Competing Interest Statement

The authors have declared no competing interest.

